# Spatiotemporal profiling reveals the impact of caloric restriction on mammalian brain aging

**DOI:** 10.1101/2025.05.04.652093

**Authors:** Zehao Zhang, Alexander Epstein, Chloe Schaefer, Abdulraouf Abdulraouf, Weirong Jiang, Wei Zhou, Junyue Cao

## Abstract

Aging induces functional declines in the mammalian brain, increasing its vulnerability to cognitive impairments and neurodegenerative disorders. Among various interventions to slow the aging process, caloric restriction (CR) has consistently demonstrated the ability to extend lifespan and enhance brain function across different species. Yet the precise molecular and cellular mechanisms by which CR benefits the aging brain remain elusive, especially at region-specific and cell type-specific resolution. In this study, we performed spatiotemporal profiling of mouse brains to elucidate the detailed mechanisms driving the anti-aging effects of CR. Utilizing highly scalable single-nucleus genomics and spatial transcriptomics platforms, *EasySci* and *IRISeq*, we profiled over 500,000 cells from 36 mouse brains across three age groups and conducted spatial transcriptomic analysis on twelve brain sections from aged mice under CR and control conditions. This comprehensive approach allowed us to explore the impact of CR on over 300 cellular states and assess region-specific molecular alterations. Our findings reveal that CR effectively modulates key aging-associated changes, notably by delaying the expansion of inflammatory cell populations and preserving cells critical to the neurovascular system and myelination pathways. Moreover, CR significantly reduced the expression of aging-associated genes involved in oxidative stress, unfolded protein stress, and DNA damage stress across various cell types and regions. A notable reduction in senescence-associated genes and restoration of circadian rhythm genes were observed, particularly in ventricles and white matter. Furthermore, CR exhibited region-specific restoration in genes linked to cognitive function and myelin maintenance, underscoring its targeted effects on brain aging. In summary, the integration of single-nucleus and spatial genomics provides a novel framework for understanding the complex effects of anti-aging interventions at the cellular and molecular levels, offering potential therapeutic targets for aging and neurodegenerative diseases.

## Introduction

Aging induces functional decline in the mammalian brain, increasing its vulnerability to cognitive impairments and neurodegenerative disorders^1^ - a process driven by alterations in the highly heterogeneous populations of brain cells. Among the various interventions to slow aging and delay age-related diseases, caloric restriction (CR) is particularly notable for its consistency in extending lifespan across species including worms, flies, rats, and mice^2–5^. Importantly, CR has demonstrated beneficial effects on brain function, enhancing learning and memory, and increasing resilience against neurodegenerative diseases^6^. These findings suggest that CR could mitigate critical molecular and cellular changes associated with brain aging. However, the intricate diversity of brain cell populations and anatomical regions limits the potential of bulk-level molecular studies to elucidate how CR affects specific cell populations and regions. As a result, the exact molecular and cellular mechanisms by which CR benefits the aging brain remain elusive.

Recent advances in single-cell and spatial transcriptomics have enabled precise measurement of gene expression changes across distinct cell populations and brain regions^7,8^. However, the low throughput of current approaches remains a challenge^6,9^, impeding our ability to comprehensively analyze the dynamics of hundreds of different brain cell states in response to anti-aging treatments, particularly for rare yet critical aging-associated cell populations (*e.g.,* neurogenic cells and activated microglia). To address the throughput limitations in single-cell and spatial genomics, we have recently developed two scalable approaches, *EasySci*^10^ and *IRISeq*^11^, which enable comprehensive single-cell and spatial transcriptomic analysis of the mammalian brain across ages and conditions.

In this study, we first applied the *EasySci*^10^ approach to analyze the transcriptional profiles of more than 500,000 nuclei derived from 36 individual mouse brains, spanning different ages and under CR or *ad libitum* (AL) feeding. This analysis enabled us to identify specific genes and pathways influenced by CR in various cell populations. Additionally, we pinpointed aging-associated cell population dynamics that can be restored by CR and the underlying genetic mechanisms. Next, we utilized *IRISeq*^11^, an optics-free spatial transcriptomics approach, to investigate the molecular changes in different brain regions upon CR. Specifically, we analyzed spatial expression profiles across twelve brain sections from aged mice with and without CR. This revealed both widespread and region-specific alterations in gene expression following CR, offering an in-depth view of the molecular and cellular pathways through which CR exerts beneficial effects on the aging brain. Overall, our spatiotemporal genomic analysis of the mouse brain offers new insights into the intricate regulatory mechanisms of CR across diverse cellular populations and brain regions, identifying potential therapeutic targets that mediate the beneficial effects of caloric restriction.

## Results

### Single-nucleus transcriptomic profiling of brain cell dynamics in response to caloric restriction

To achieve a comprehensive understanding of how various brain cell populations and regions respond to caloric restriction (CR), we applied two innovative approaches, *EasySci*^10^ and *IRISeq*^11^, which we recently developed for scalable single-nucleus and spatial genomic analysis of the mammalian brain across different ages and conditions. We first conducted single-nucleus transcriptomic analysis on thirty-six mouse brain samples from a diverse cohort spanning three age groups: 12, 18, and 24 months (**Figure 1A, upper**). Each age group includes 12 individual brains, equally divided between those undergoing 60% caloric restriction initiated at 4 months of age and *ad libitum-fed* controls (**Table S1**). As expected, CR significantly reduced the body weight of animals in all age groups, while its impact on brain weight was relatively minor (**Figure 1B, Figure S1A**). In addition, we applied *IRISeq* to assess region-specific gene expression dynamics. This analysis covered twelve brain sections from six aged (24-month-old) mice in CR or control conditions, yielding a total of 65,128 spatially barcoded transcriptome profiles (**Figure 1A, lower; Figure 1D**).

**Figure 1.**
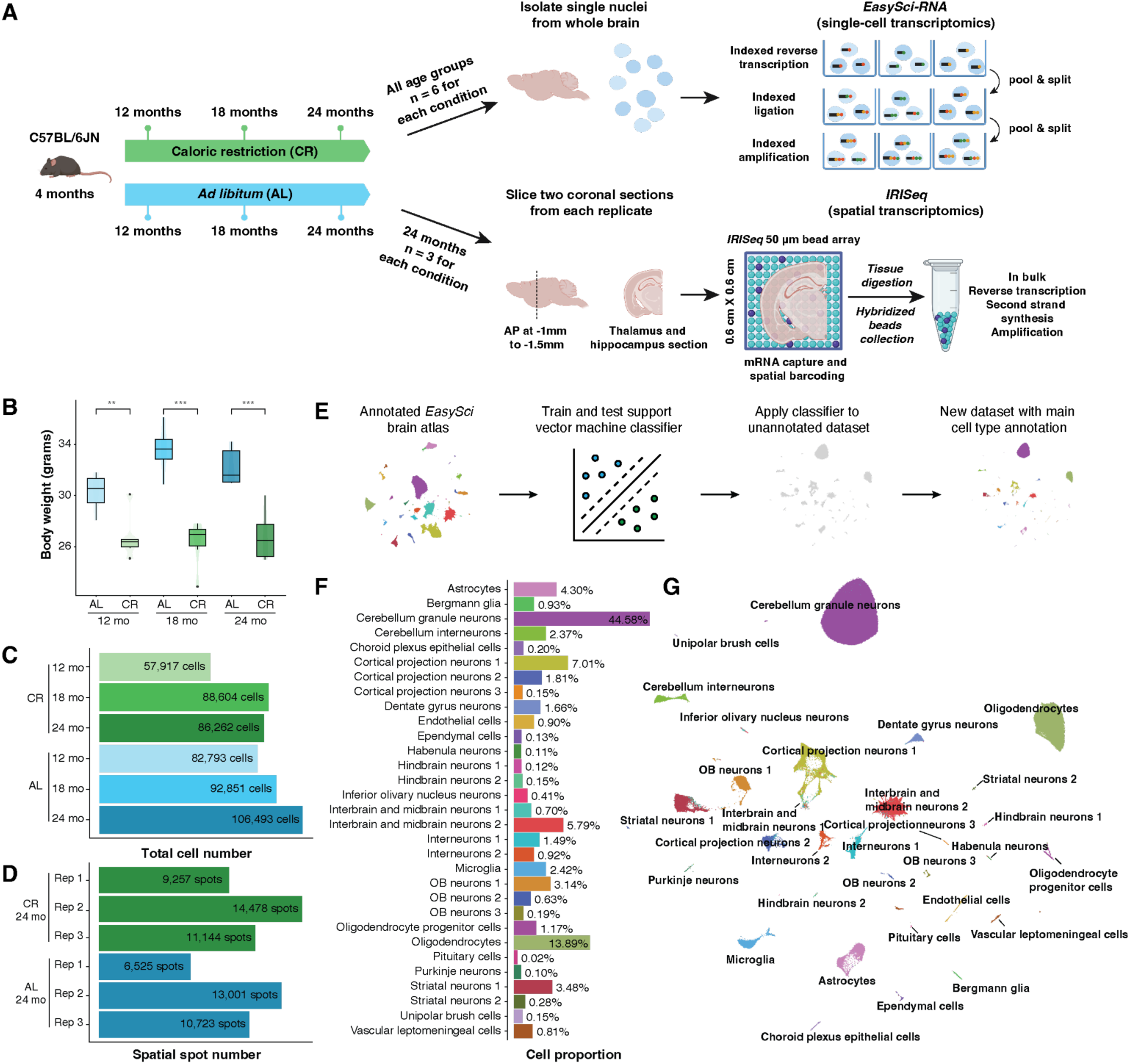
Single-nucleus and spatial transcriptome profiling of mouse brains across various ages and in response to CR. **(A)** The scheme of experimental design. Key steps are outlined in the texts. AP, anterior-posterior coordinates. **(B)** Body weights across experimental groups. *Welch’s t-test*,** represents p-value < 0.01, *** represents p-value < 0.001, and ‘NS.’ represents not significant. **(C)** Barplot showing total nucleus numbers recovered per experimental group in the *EasySci* experiment. **(D)** Barplot showing spatial spot numbers recovered per replicate in the *IRISeq* experiment. **(E)** Schematic representation of the key steps involved in the annotation of main cell types. **(F)** Barplot showing the fraction of each main cell type from single-cell RNAseq data, normalized by the total number of cells. **(G)** Uniform Manifold Approximation and Projection (UMAP) visualization of current single-cell study (referred to as “CR brain atlas”), colored by main cell type annotation.

Following sequencing of the *EasySci* library and filtering out low-quality cells and doublets, we obtained 514,920 high-quality single-nucleus gene expression profiles across various conditions. The number of nuclei profiled per age and diet condition ranged from 57,917 to 106,493, with an average of 14,303 nuclei recovered per individual (**Figure 1C, Figure S1B**). Each single-nucleus profile contained an average of 5,369 unique transcripts (UMIs) and 1,504 genes (and a median of 1,994 UMIs and 974 genes) (**Figure S1D**). Notably, all samples were processed and sequenced concurrently in the same experiment, resulting in minimal experimental batch effects (**Figure S1E-F**).

Next, we applied a machine learning approach to automate annotation of the main cell types recovered in the current single-nucleus study (referred to as “*CR brain atlas*”), by transferring labels from a previously published single-cell mouse brain aging atlas^10^ (referred to as “*reference brain atlas*”). Specifically, we trained a support vector machine-based classifier using data from the reference brain atlas and then applied it to annotate each single-nucleus transcriptome profiled in this study (**Figure 1E**). We performed 5-fold cross-validation in the reference dataset to validate classifier accuracy, obtaining substantially higher F1-score (**Figure S2 A-D**). We identified all 31 main cell types defined in reference brain atlas^10^ (**Figure 1F-G**), including abundant cell types such as cerebellum granule neurons (44.58%) and oligodendrocytes (13.89%), as well as rare cell types like Purkinje neurons (0.10%) and pituitary cells (0.02%). Unsupervised dimensionality reduction applied to the CR brain atlas data also yielded highly distinct cellular clusters that closely matched the annotated cell types (**Figure S1C)**, validating our annotation method. The cell annotations were further validated by integrating our data with the reference brain atlas^10^, which demonstrated a high degree of consistency of the same cell types (**Figure S1G**). Although cell type proportion information was not used during label transfer, we observed significant correlations in main cell proportions between both the query and reference datasets (Spearman’s rho = 0.85, p-value = 5.0e-07) (**Figure S1H**). The distinct molecular states of these cells were further confirmed by cell-type-specific gene signatures (**Figure S2E**), such as choroid plexus epithelial cells marked by *Ttr*^12^, vascular leptomeningeal cells marked by *Slc47a1*^13^, and microglia marked by *Csf1r*^13^. In addition to our primary annotation, we also provide cell type labels derived from the Allen Brain Initiative Cell Census Network (BICCN) reference, which show strong alignment with our dataset **(Fig. S1I-K)**.

### Caloric restriction rescues age-related gene expression changes

To assess the impact of aging and calorie restriction (CR) on brain gene expression, we first aggregated single-cell transcriptomes from individual brains and performed principal component analysis (PCA) (**Figure S3A-B**). The *ad libitum* samples exhibited a smooth trajectory from 12 to 18 to 24 months, reflecting a consistent age-related progression (**Figure S3A**). Although CR did not markedly alter the bulk brain transcriptomes at 12 and 18 months, at 24 months the CR-treated brains shifted significantly toward a younger transcriptional state (p-value = 0.014), albeit with higher variability among individual samples compared with the *ad libitum* controls (**Figure S3 A-B**).

To address the variation in CR samples at 18 and 24 months and to enhance statistical power, we combined these two time points for differential expression gene analysis. Using stringent criteria - an FDR of 10e^-5^ and at least a two-fold change between adult (12 months) and aged (18 and 24 months combined) brains - we identified 456 cell-type-specific DE genes (**Table S2**). This combined approach not only showed strong concordance with DEGs identified in the direct 12-month vs. 24-month comparison but also yielded more robust results (**Figure S3C**). Notably, certain cell types—vascular leptomeningeal cells, olfactory bulb (OB) neurons, and inferior olivary nucleus neurons—exhibited a greater number of DE genes, suggesting significant molecular alterations during aging (**Figure 2A**).

**Figure 2.**
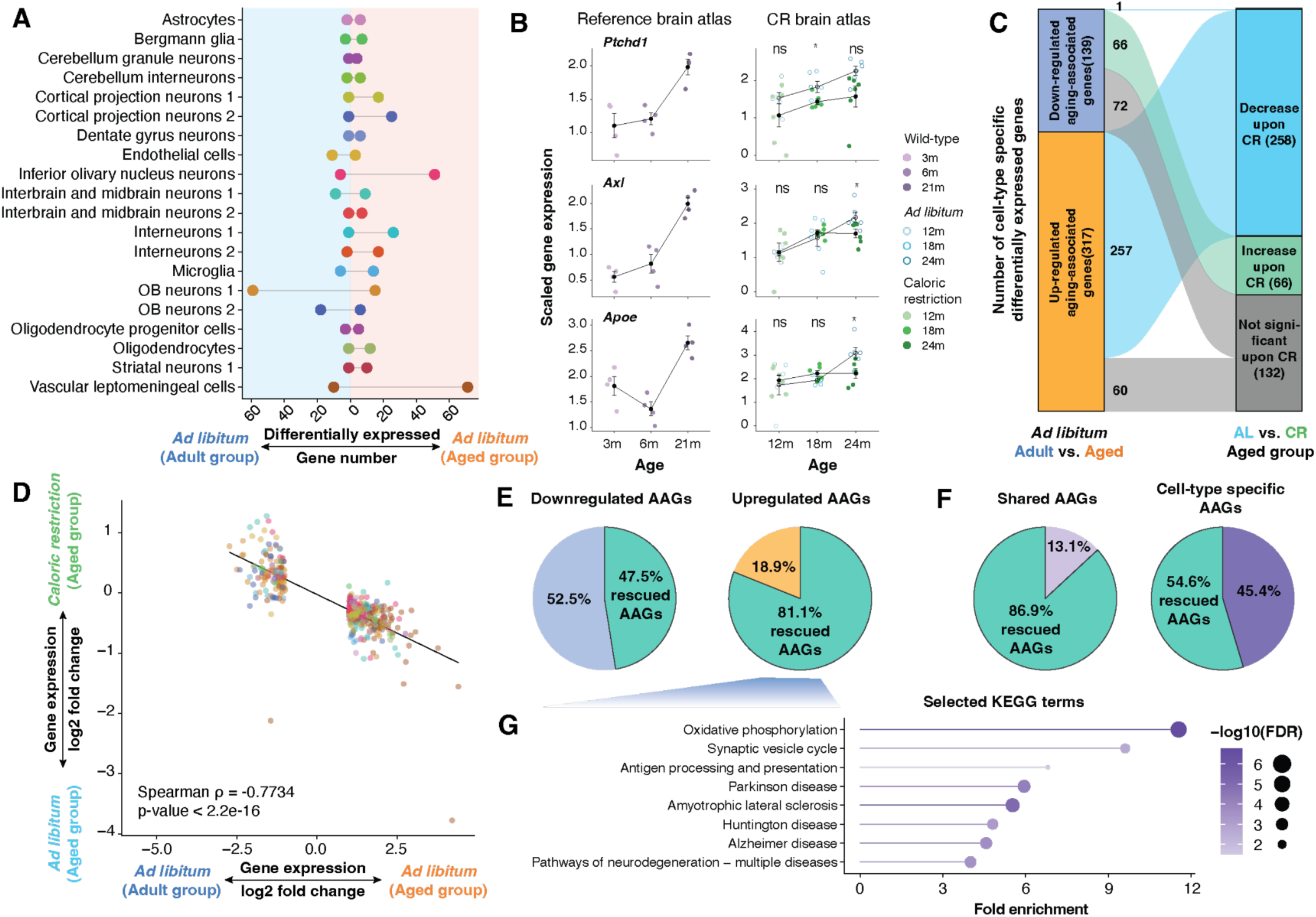
Caloric restriction reverses aging-associated gene expression changes. **(A)** Barplot showing the number of differentially expressed genes between adult (12 months) and aged (18 months and 24 months) mice across main brain cell types under *ad libitum* condition. Cell types with at least 1,000 nuclei in the AL dataset are shown. **(B)** Line plots showing the aggregated expression for each replicate of selected AAGs in microglia across age groups and conditions in the published *EasySci* brain aging atlas^10^ (Left) and this study (Right). Colors indicate dataset and diet groups. Welch’s *t*-test, ** represents p-value < 0.01, * represents p-value < 0.05, and ‘ns’ represents not significant. **(C)** Sankey diagram showing the rescuing effect of caloric restriction on cell-type-specific AAGs. Cell-type-specific AAGs, identified by comparing adult and aged groups under *ad libitum* conditions for each main cell type, are shown on the left. The impact of CR is evaluated by comparing AL and CR nuclei within the aged group. Genes are defined as “rescued” if they are upregulated with aging and significantly decreased under CR, or downregulated with aging and significantly increased under CR (FDR of 0.05). Flow widths represent the number of cell-type-specific genes in each category. **(D)** Scatter plots comparing the alterations in differentially expressed genes between CR vs. AL and aged vs adult, including a linear regression line. Each gene is colored by the main cell type it comes from and only AAGs are shown. **(E-F)** Pie charts depicting the proportions of AAGs rescued by CR across different categories: E (left) shows down-regulated AAGs, E (right) shows up-regulated AAGs, F (left) highlights AAGs shared across more than one cell type, and F (right) displays AAGs unique to specific cell type. **(G)** Lollipop plot showing the top enriched Kyoto Encyclopedia of Genes and Genomes (KEGG) terms for the genes that are significantly increased in aging and rescued by CR.

Most of these DE genes were specific to individual cell types (**Figure S3D**). For example, aged microglia showed increased expression of genes linked to cellular proliferation (*e.g., Ptchd1*^14^), and genes involved in motility and phagocytic activity (*e.g., Myo1e*^15^), accompanied by increased levels of genes associated with lipid and amyloid-beta metabolism (*e.g., Apoe*^16^), and immune response (*e.g., Axl*^17^). These cell-type-specific gene expression changes were further confirmed by previous *EasySci* brain atlas^10^, suggesting dysregulated microglial activation in the aged brain (**Figure 2B**). Meanwhile, we identified 49 genes that are significantly altered across multiple cell types with a highly consistent trend (**Figure S3D**). These genes were involved in pathways such as intercellular communications (*e.g., Rab3a, Scg2*), genomic instability (*e.g., Tspy14, Oip5os1*), and loss of proteostasis (*e.g., Hsp90aa1, Hsp90ab1*) (**Figure S3D-E**).

We next examined the impact of CR on rescuing age-related changes in gene expression. A substantial fraction (324 out of 456) of cell-type-specific age-associated gene signatures (AAGs) showed significant changes between CR and *ad libitum* conditions at 24 months (FDR of 0.05) (**Figure 2C**). Remarkably, nearly all of the cell-type-specific genes that were significantly changed in both aging and CR (323 out of 324) were changed in opposite directions in aging and CR (**Figure 2C-D**), suggesting CR can effectively mitigate age-related changes in gene expression. Furthermore, while some genes (*e.g., Ptchd1* in microglia) are rescued by CR at an earlier life stage (*i.e.,* 18 months), others, like *Apoe* and *Axl* in microglia, only show significant differences between CR and control conditions at a later life stage (*i.e.,* 24-month) (**Figure 2B**). This differential response likely results from elevated cellular stress in later life, underscoring the age-dependent effects of CR and highlighting the importance of including multiple time points to evaluate the effectiveness of anti-aging interventions.

Interestingly, CR does not uniformly rescue aging-associated genes. For instance, 81.1% of genes up-regulated in aging are rescued by CR, whereas only 47.5% of down-regulated genes can be rescued (**Figure 2E**). Genes that increase with aging but are mitigated by CR are enriched in mitochondrial functions related to oxidative phosphorylation, immune signaling, antigen processing, and neurodegeneration processes (**Figure 2G**, **Figure S3F**). Furthermore, genes common across multiple cell types are more likely to be rescued by CR. For example, 86.9% of shared aging-associated genes are significantly rescued by CR, compared to only 54.6% of cell-type-specific genes (**Figure 2F**). This difference is partially attributed to the predominance of up-regulated genes among the shared genes. However, even when focusing exclusively on up-regulated genes, shared aging-associated genes are more frequently rescued by CR (90%) than those unique to specific cell types (67%). This suggests that CR preferentially mitigates widespread increases in gene expression, such as those related to cellular stress responses, rather than targeting genes specific to individual cell types (**Figure S3G)**.

### The impact of caloric restriction on reversing age-related cell population changes

Caloric restriction mitigates many aging-associated gene expression changes, either by uniformly reversing these changes across all cells of a given type or by modulating the abundance of specific subpopulations enriched for age-related molecular signatures. To distinguish between these two mechanisms, we investigated the impact of CR on the cell abundance dynamics of aging-associated cell subpopulations. Similar to the computation pipeline for annotating main cell types, we trained a support vector machine-based classifier that utilized the same reference brain atlas^10^ to annotate the detailed molecular state of each cell in this study (**Figure 3A, Figure S2D**). In total, we annotated over 300 subpopulations with distinct molecular profiles, many of which represent either transitional states within a broader cell type (e.g., disease-associated microglia) or rare cell types that form discrete clusters upon sub-clustering (e.g., T cells and B cells). While our label transfer and cellular state annotation algorithm do not provide cellular population counts, there is a significant correlation in the population distributions of subpopulations between the two datasets (Spearman rho = 0.76, p-value < 2.2e-16, **Figure 3B**), demonstrating that our algorithm preserves cellular population proportions.

**Figure 3.**
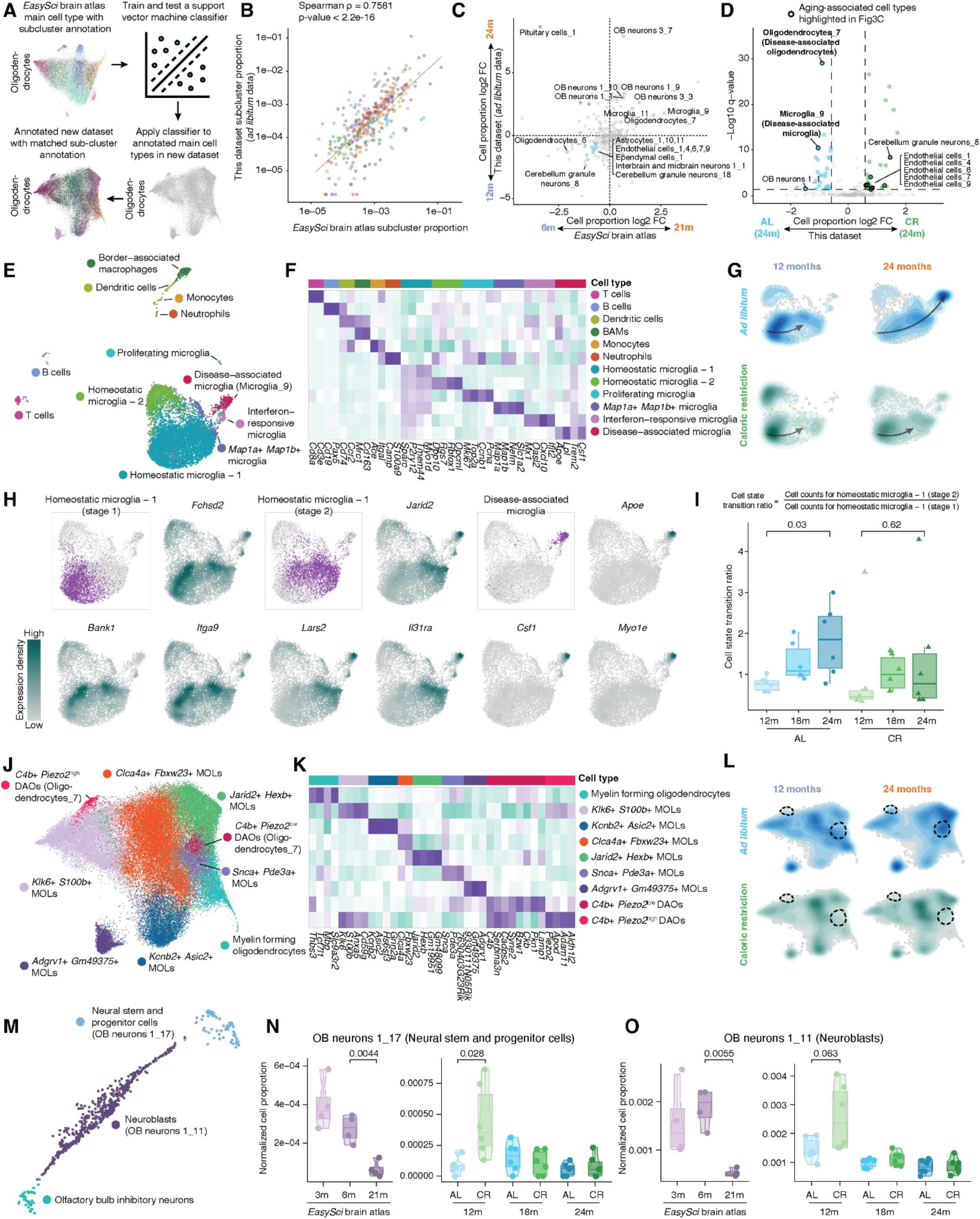
Caloric restriction rescues aging-associated cell population dynamics. **(A)** Diagram depicting the computational pipeline to transfer the subpopulation labels from the reference brain aging atlas^10^ to this dataset. **(B)** Scatter plot showing the correlation of overall subpopulation proportions between the reference brain aging atlas^10^ and this CR brain atlas study (*ad libitum* data only). **(C)** Scatter plot showing the correlation of aging-associated subpopulation proportion changes between the reference brain aging atlas^10^ (6 months vs. 21 months) and this study (12 months vs. 24 months, *ad libitum* data only). **(D)** Volcano plot showing the cell subpopulation populations significantly altered between caloric restriction and *ad libitum* animals at 24 months. Aging-associated cell populations that exhibit similar aging dynamics between the reference brain aging atlas^10^ and this dataset (*ad libitum*) were highlighted in a black circle. **(E)** UMAP visualization of microglia by integrating cells from this study and two external datasets^10,19^, colored by subpopulation annotation. **(F)** Heatmap showing the marker gene expressed in each subpopulation identified in **(E)**. **(G)** UMAP Density plot showing the cellular distribution of microglia over the progress of aging in CR and control conditions. The arrow indicates the inferred cell state transition trajectory. **(H)** UMAP visualization of microglia cell states mapping and corresponding molecular markers. **(I)** Box plot detailing the proportion change between two homeostatic microglia cell states during aging and caloric restriction. **(J)** UMAP visualization of oligodendrocytes by integrating cells from this study and two external datasets^10,19^, colored by subpopulation annotation. **(K)** Heatmap showing the marker gene expressed in each subpopulation identified in **(H)**. **(L)** Density plot showing the distribution of oligodendrocytes in adult and aged mice from CR and control conditions. Two disease-associated oligodendrocyte states are highlighted. **(M)** UMAP visualization of cells involved in neurogenesis by integrating cells from this study and two external datasets^10,19^, colored by subpopulation annotation. **(N-O)** Box plots showing the cell proportion changes in aging-depleted neural stem and progenitor cells (N) and neuroblasts (O) across different ages and diet conditions in the brain aging atlas^10^ (Left) and this dataset (Right).

Once subpopulations were annotated using our data-driven approach, we performed differential abundance (DA) analysis to pinpoint cell subpopulations significantly altered with aging in the CR brain atlas, and compared these results with those from reference aging brain atlas. Despite the different time frames of our study (12 months vs. 24 months) and the reference brain atlas^10^ (6 months vs. 21 months), 21 out of 25 aging-associated subpopulations identified in both datasets (FDR of 0.05, at least 1.5 fold change between adult and aged brains) exhibit consistent aging-associated dynamics (**Figure 3C**). The expanded cell subpopulations include disease-associated microglia (*Microglia-9*, marked by *Csf1* and *Apoe*), disease-associated oligodendrocytes (*Oligodendrocyte-7*, marked by *C4b* and *Serpina3n*), and several OB neuron subpopulations. The aging-associated expansion of these glial cells aligns with prior single-nucleus and spatial genomic studies^10,11^, suggesting increased inflammatory stress in the aged brain. Conversely, we observed depletion in several endothelial cell subpopulations (**Figure 3C**), indicating damage to the neurovascular system and compromised blood-brain barrier function in the aged brain. Additionally, we detected a reduction in myelin-forming oligodendrocytes (Oligodendrocyte-6, marked by *Tcf7l1 and Slc9a3r2*), consistent with the diminished remyelination capacity observed in the aged brain^18^.

We next examined the effects of CR on reversing age-related cell population changes. Notably, CR significantly reversed changes in a substantial number of aging-associated cell subpopulations (12 of the 21; FDR of 0.05 with over 1.5 fold changes between CR and control) (**Figure 3D**). For example, several endothelial subpopulations, which typically deplete at 24 months, were restored following CR(**Figure 3D**). This observation is aligned with previous studies suggesting that CR helps maintain cerebral vascular homeostasis^6^. In addition, CR effectively mitigated the aging-associated expansion of several reactive glial cell populations, including disease-associated microglia (Microglia-9) and disease-associated oligodendrocytes (Oligodendrocyte-7) (**Figure 3D**, **Figure S4A, E**), indicating a reduction in brain inflammatory stress upon CR.

To delve deeper into the detailed cell state transitions underpinning the effect of CR, we focused on several glia subpopulations where significant rescue effects of CR were noted. We first integrated single-nucleus transcriptomes of microglia from our dataset with those profiled in the reference brain aging atlas^10^ and a mouse progenitor cell atlas^19^ (**Figure 3E)**. This integration identified twelve distinct cellular states, including various immune cell populations (*e.g.,* T cells, B cells, dendritic cells) and several other microglia subpopulations (*e.g.,* homeostatic microglia, proliferating microglia, *Cxcl10+* interferon-responsive microglia) (**Figure 3E)**. The distinct molecular states of these clusters were further confirmed by validation across three independent datasets (**Figure S4B)**, and the expression of cluster-specific gene signatures (**Figure 3F, Figure S4C-D)**. Consistent with our cell population analysis, we observed an age-dependent transition from a homeostatic microglia state to disease-associated microglia (DAM) state, and CR can effectively inhibit the expansion of DAM by obstructing this transition **(Figure 3G).** Additionally, we identified a less-characterized microglia state (*Jarid2*+, *Lars2*+, *Il31ra*+) that exhibits characteristics of both homeostatic microglia and DAM, indicating an intermediate state between the two (**Figure 3H**). Notably, aging facilitates the transition into this *Il31ra*+ *Jarid2*+ intermediate microglia state, a process that can be effectively mitigated by CR(**Figure 3I**).

Similarly, we delved into oligodendrocyte dynamics by integrating single-nucleus transcriptomes of oligodendrocytes with two external datasets^10,19^. We revealed detailed oligodendrocyte subpopulations (*e.g.,* disease-associated oligodendrocyte ‘DAO’ subpopulations, mature oligodendrocyte ‘MOL’ subpopulations) and associated molecular signatures (**Figure 3J-K, Figure S4E-H)**. Notably, we observed two *C4b*+ disease-associated oligodendrocyte populations (*C4b*+ *Piezo2*^high^ DAOs and *C4b*+ *Piezo2*^low^ DAOs) that transit from distinct oligodendrocyte subpopulations, which were both reversed with CR (**Figure 3L)**.

Depleted neurogenesis is a hallmark of brain aging. To examine the effects of CR on adult neurogenesis, we integrated neurogenic cells (neural stem and progenitor cells and neuroblasts) from our study with neurogenic cells recovered from the progenitor cell atlas^19^. This integration allowed us to delineate a continuous cellular differentiation trajectory from progenitor cells to OB neuroblasts (**Figure 3M, Figure S5A-B**), aligning with findings from our previous studies^19^. Aging typically triggers a rapid depletion of neurogenic cells at an early stage^20^, which is observable in our reference EasySci brain dataset **(Figure 3N)**. In our CR dataset, we did not observe a further depletion of neuronal stem and progenitor cells or OB neuroblasts in AL samples profiled after 12 months (**Figure 3N-O**), suggesting that neurogenic cells may already be depleted at this point. In contrast, the CR samples exhibited an average of 1.8-fold increase in neural stem and progenitor cells and a 5-fold increase in neuroblasts at 12 months (**Figure 3N-O**). Further differential expression analysis was performed on OB neurogenic cells from 12-month-old mice to identify genes regulated by CR (**Fig. S5C-D**). We identified 57 genes upregulated and 34 genes downregulated under CR versus AL. Upregulated genes (*e.g., Top2a, Egfr*) were associated with enhanced proliferation, while downregulated genes related to neuronal differentiation processes (*e.g., cilium assembly*) and negative regulation of canonical Wnt signaling (**Fig. S5E-F**), suggesting that CR can expand the OB progenitor pool by promoting proliferation and suppressing neuronal differentiation. However, CR did not alleviate the depletion of neurogenesis at later stages (18 and 24 months) (**Figure 3N-O**). This indicates that while CR delays the depletion of neurogenic cells beyond the 12-month timepoint, it does not completely prevent it, underscoring the importance of including multiple time points in our study to capture the age-dependent effects of CR on cell population dynamics.

### Spatial transcriptome profiling reveals region-specific gene targets of caloric restriction

Several aging-associated cell population changes rescued by caloric restriction, including depletion of neurogenic cells and expansion of disease-associated microglia (DAM) and disease-associated oligodendrocytes (DAOs), occur in populations enriched in ventricles and white matter, suggesting a significant rescue impact of CR in these regions. To validate this effect and pinpoint region-specific molecular responses to CR, we utilized a scalable spatial transcriptomics platform, *IRISeq*^11^, to profile twelve brain sections from six 24-month-old mice under both CR and AL conditions (**Figure 1A**). We sectioned the brain to include regions such as the cortex, hippocampus, midbrain, hypothalamus, and white matter areas, all of which exhibit significant structural and functional changes during aging^21^. The *IRISeq* platform employs 50 µm barcoded gel beads to capture the spatial distribution of gene expression across different brain regions, reconstructing spatial interactions based on bead-bead interaction oligos^11^. With this platform, we recovered a total of 65,128 transcriptionally barcoded spatial spots across twelve brain sections, detecting a median of 2,700 unique transcripts per spot (**Figure 4A**).

**Figure 4.**
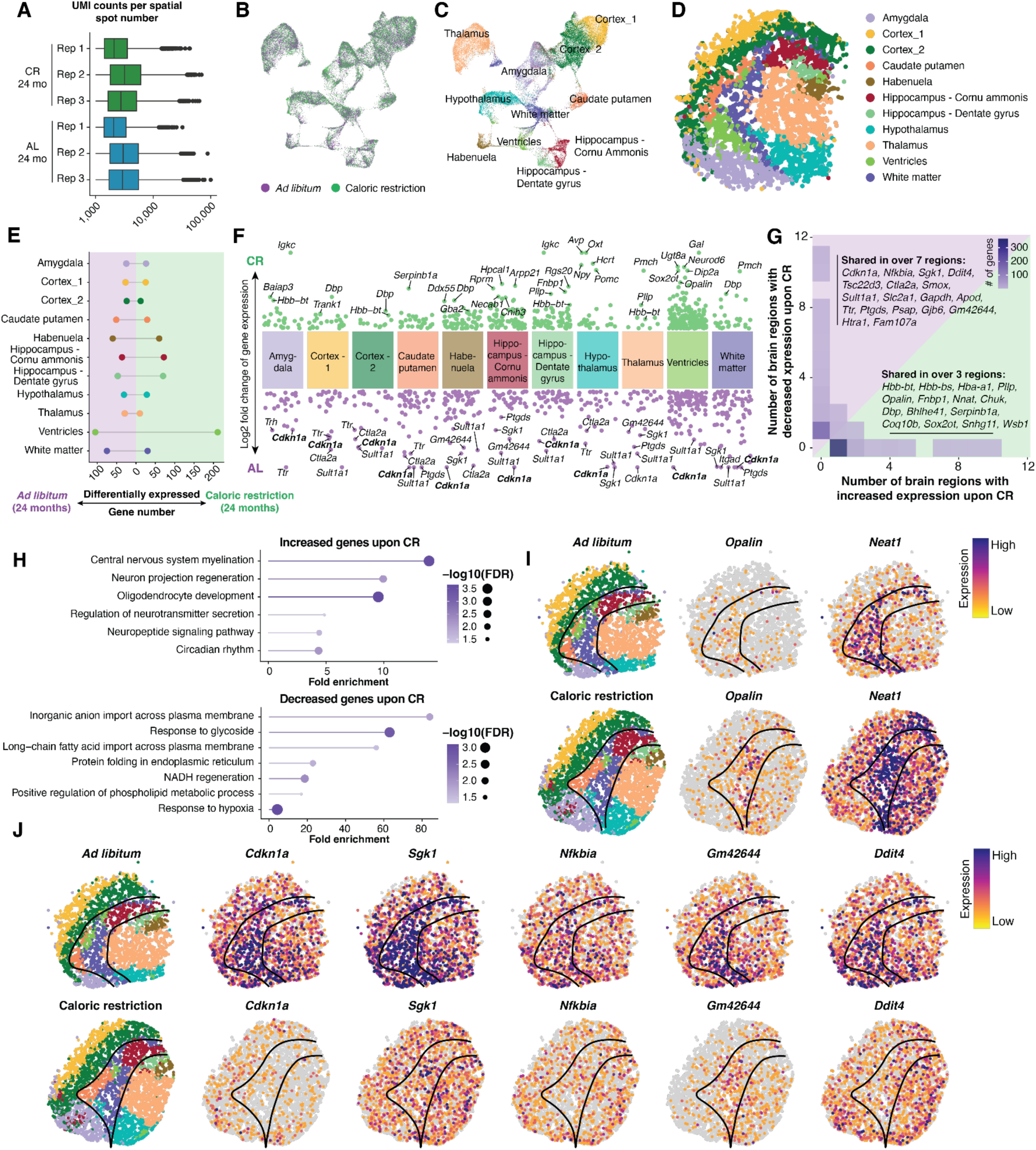
The region-specific molecular responses to caloric restriction. **(A)** Boxplot showing the number of unique transcripts detected per ‘spatial spot’ across brain sections for each mouse individual. **(B-C)** UMAP plot showing the gene expression clusters of spatial spots, colored by dietary conditions (B) and annotated brain regions (C). **(D)** Reconstructed image of all spatial spots derived from bead-bead connections in the *IRISeq* profiling, with each spot colored by annotated brain regions. **(E)** Barplot showing the number of CR-associated DE genes across various brain regions, colored by upregulated (green) and downregulated (purple) genes upon CR. **(F)** Dot plot showing the differentially expressed genes between CR and AL groups (24 months) across brain regions. Top gene markers enriched in each diet condition are annotated. **(G)** Heatmap showing the number of brain regions where genes are either upregulated (x-axis) or downregulated (y-axis) in response to CR. The color gradient from light to dark blue indicates the number of genes, with darker shades representing higher numbers. **(H)** Lollipop plot showing the top enriched GO Biological Process terms for the CR-associated genes that are up-regulated (upper) or down-regulated (lower). **(I-J)** Reconstructed spatial maps display the expression levels of upregulated (I) and downregulated (J) genes in response to CR. The raw gene expression count per spot is normalized by the total count, multiplied by 10,000, and log-transformed.

To annotate spatially barcoded transcriptome profiles across brain regions, we integrated transcriptome data from all twelve sections for clustering analysis, which identified eleven transcriptionally distinct clusters corresponding to specific brain regions (**Figure 4B-C**). These clusters were then annotated based on region-specific gene features (**Figure S6A-B**, **Table S3**). Using bead connection data, we reconstructed the spatial distribution of each receiver bead across different sectioning regions (**Figure 4D**). This reconstruction accurately mapped the receiver beads to specific brain regions, consistent with their gene expression-based annotations (**Figure 4D, Figure S6C**). The top region-specific gene features identified are highly consistent with previous studies, highlighting markers such as *Plp1* for white matter, *Pmch* for the hypothalamus, and *Camk2n1* for the cortex^11^ (**Figure S6C**). These results underscore the capacity of the *IRISeq* platform to perform spatial transcriptome profiling without the need for pre-indexed oligo arrays or optical imaging.

We next investigated region-specific gene expression changes in response to caloric restriction. Through differential expression analysis, we identified 700 DE genes that are significantly altered between CR and AL groups across various regions. Notably, the ventricle region displayed the highest number of DE genes (**Figure 4E**), aligning with the significant rescue effect of CR on cell populations (*e.g.,* neuronal progenitor and glial cells) in this area. Most of these CR-associated DE genes (559 out of 700) exhibit high region specificity (**Figure 4F-G**). For example, in the hypothalamus, the top five genes significantly upregulated by CR represent activation of specific neuronal populations closely associated with calorie regulation and metabolic control. Notably, these include *Avp* (75-fold increase), linked to cognitive function and known to suppress food intake when activated^22^; *Oxt* (34-fold increase), a hypothalamic neurohormone implicated in social behavior and appetite suppression in both humans and animal models^23^; *Hcrt* (orexin/hypocretin), involved in sleep behavior regulation and neuronal activation by dietary amino acids^24,25^; *Pomc*, associated with promoting satiety and reducing appetite^26^; and *Npy*, known for its crucial role in energy homeostasis and significantly induced in the hypothalamus under caloric restriction (**Figure S6D-H)**^27^. Additionally, CR elevated the expression of oligodendrocyte-associated genes, such as *Ugt8*, *Sox2ot*, *Aspa*, and *Dip2a* (**Figure 4F**), with prominent upregulation in the ventricular region, supporting a beneficial role of CR in oligodendrocyte maturation and maintenance in the aging brain. These findings were further corroborated by analyses of spatial transcriptomic data across biological replicates, reinforcing the robustness of our observations. (**Figure S6D-H).**

Meanwhile, 141 out of 700 CR-associated genes are upregulated across more than one brain region, with most exhibiting highly consistent changes across these areas (**Figure 4G**). We witnessed a widespread upregulation of circadian rhythm genes (*e.g., Dbp*^28^, *Bhlhe41*^29^) in response to caloric restriction (**Figure 4G-H**), suggesting that caloric restriction contributes to restoring the dysregulated circadian rhythms observed in the aged brain^30^. This effect is complemented by the CR-induced upregulation of genes in the neuropeptide signaling pathway (*e.g., Pmch*^31^), which are crucial for regulating circadian and feeding behaviors (**Figure 4F-G**). Additionally, CR enhances the expression of genes associated with myelination (*e.g., Pllp*^32^, *Opalin*^33^) across the ventricles, white matter, and hippocampus regions (**Figure 4F, H-I**), which is consistent with previous findings that caloric restriction can ameliorate myelin degeneration in neurodegenerative disorders^34^, and aligned with the top enriched Gene Ontology terms with genes that are upregulated under CR conditions (**Figure 4H, upper**).

In addition, we identified 68 genes that are significantly depleted across more than one brain region in response to calorie restriction. Notably, many of these CR-depleted genes are key players in senescence pathways, including *Cdkn1a*, *Cdkn1c*, *Bcl2l1*, and *Igfbp7* (**Figure 4F-G, Figure S6I-L**). The reduction in the expression of these senescence-associated genes in the ventricles and white matter is correlated with decreased cellular stress in these regions, evidenced by the suppression of genes involved in DNA damage (*e.g., Ddit4*, *Gadd45g*), inflammation (*e.g., Nfkbia*, *A2m*, *Tsc22d3*), and oxidative stress (*e.g., Sgk1*, *Txnip*) (**Figure 4F-G, J**). To validate CR’s effect on reducing senescence, we employed the SenMayo gene set^35^—a curated panel of 125 validated genes associated with cellular senescence—to assess major cell types in our single-cell dataset. Our analysis showed that for most cell types, SenMayo scores increased with age and decreased under CR **(Figure S7A**), with aging effects generally stronger than dietary effects. Notably, in microglia and oligodendrocytes, both age-related increases and CR-induced decreases were significant after FDR correction (**Figure S7B**). In oligodendrocytes, senescence score increased across several subclusters with increasing age **(Figure S7C)**, with no subcluster driving the phenomenon. By contrast, the rise in microglia SenMayo scores was largely driven by the accumulation of microglia-9 (DAM^36^), which exhibited the highest SenMayo score of any subcluster in 24-month-old *ad libitum*-fed mice (**Figure S7D**). These results are consistent with previous studies^37^ that identified senescent microglia as either a subset of DAM or as a separate signature expressing markers such as *Trem2* and *Apoe*; in our study, we suspect that the limited number of microglia prevented clear discrimination between senescent microglia and DAM. Overall, these transcriptomic analyses further support our spatial genomic results, confirming that CR significantly modulates cellular senescence across brain cell types.

Additionally, CR significantly alters the region-specific expression of lesser-characterized non-coding RNAs, notably increasing *Neat1* and depleting *Gm42644* (**Figure 4I-J**). These changes, including the reduction of senescence markers and cellular stress signals, along with the upregulation of myelination pathways, are predominantly observed in the ventricles and white matter of the aged brain (**Figure 4I-J**). Such alterations align with the impact of caloric restriction on reversing age-related cell population changes (*e.g.,* decreased disease-associated oligodendrocytes and disease-associated microglia) in the same regions, underscoring its potential therapeutic implication for the functions of these critical brain regions in both aging and neurodegenerative diseases.

## Discussion

While many anti-aging strategies have shown promise in mitigating the functional decline associated with aging in the mammalian brain^38^, a comprehensive understanding of their cellular and molecular mechanisms remains elusive. This knowledge gap is largely due to the limitations of traditional methodologies, which are unable to decipher the impact of anti-aging interventions on the dynamics of hundreds of brain cell populations, particularly rare, yet crucial, aging-associated cell populations. To address this limitation, our study first applied *EasySci*^10^, an optimized single-cell/nucleus RNA-seq by combinatorial indexing strategy, to comprehensively assess the impact of calorie restriction on more than 300 cellular states by analyzing over 500,000 nuclei from 36 mouse brains at three different age stages (12, 18, and 24 months). Furthermore, we employed a recently developed scalable spatial transcriptomics platform, *IRISeq*^11^, for spatial analysis of twelve brain sections from six 24-month-old mice under both CR and control conditions. Substantial aging-related cell population changes were revealed, including the expansion of inflammatory cell states such as disease-associated microglia (DAM) and disease-associated oligodendrocytes (DAOs). We also noted an age-associated depletion of cell populations vital to neurovascular integrity (*e.g.,* endothelial cells) and the brain myelination pathway (*e.g.,* myelin-forming oligodendrocytes), consistent with previous single-cell studies^10^.

CR demonstrated a potent, cell-type-specific impact on these aging-associated cell populations. Consistent with previous studies^39^, we observed a depletion of neurogenesis-related cells (i.e., neuronal stem and progenitor cells and OB neuroblasts) before 12 months. Under CR, this decline was delayed; however, it was not entirely prevented, as evidenced by their decline to similar levels as control by 18 months. In contrast, the expansion of inflammatory cells, typically observed after 18 months, was markedly inhibited by CR during these later stages. Additionally, we noted a CR-mediated rescue effect on endothelial cell depletion at 24 months, although this was not evident at earlier stages.

To explore the molecular mechanisms underlying the benefits of calorie restriction, we analyzed cell type- and region-specific gene expression alterations in response to CR. Overall, CR significantly reduced the expression of 70.8% of aging-associated genes across various cell populations. These genes are predominantly involved in cellular stress responses (*e.g., Hsp90aa1*) and inflammatory processes (*e.g., Apod*), which typically increase brain-wide during aging. The consistent suppression of these stress responses, observed in both single-nucleus RNA sequencing and spatial transcriptomics analyses, suggests that CR effectively mitigates aging-associated cellular damage.

Previous studies have shown that accumulated cellular damage can induce cellular senescence, leading to blocked cellular proliferation and enhanced neuroinflammation^40^. In line with these findings, our study observed a global downregulation of senescence-associated genes across brain regions upon CR. This is consistent with the anti-senescence effect of CR on other mouse tissues, such as the intestine and liver^41^. Notably, the effects of CR on reversing cellular senescence and restoring cell population dynamics were particularly significant in the ventricles and white matter regions. Thus this reduction in cellular senescence likely facilitates the rejuvenation of other cell populations, such as reduced proliferation of disease-associated microglia and enhanced neurogenesis in the aged brain.

In addition to mitigating cellular stress and senescence, CR exhibited several lesser-known effects on specific cell populations and regions. For example, CR led to the depletion of specific non-coding RNAs, such as *Gm42644*, in the ventricles and white matter. Furthermore, CR upregulated several genes associated with cognitive functions and metabolic activities, such as *Avp* and *Oxt* in the hypothalamus. In the ventricles, CR significantly upregulated genes involved in neurogenesis and myelin maintenance. These changes make sense given the observed enhancement of neurogenesis following CR, underscoring the targeted effects of CR on brain function and aging.

Our analysis highlights that cell population dynamics could serve as an early indicator of mammalian aging. Unlike molecular changes, alterations in cellular populations are directly linked to the functional changes of mammalian organs. Meanwhile, it is also more robust against experimental batch effects as cellular states are defined by a combination of molecular markers. Using cell population dynamics as an early indicator, we observed that the effect of CR is apparent even at 12 months or earlier, as evidenced by the delayed depletion of neuronal stem and progenitor cells in the brain. Furthermore, we observed highly time-dependent changes in cell population dynamics, such as the depletion of neuronal stem and progenitor cells preceding the expansion of inflammatory cells. This observation aligns with our previous studies that cataloged distinct temporal dynamics of cell population changes across different age windows^42^. Such findings underscore the importance of selecting appropriate time windows for evaluating anti-aging interventions, as their effects vary significantly based on the specific cell populations they impact.

In summary, we systematically explored the impact of caloric restriction on global cell population dynamics and the associated cell type- and region-specific molecular alterations by highly scalable single-nucleus genomics and spatial genomics. Moreover, the affordability and high-throughput capabilities of these methods could be applied to decipher the cellular and molecular mechanisms for other therapeutic interventions^43^. This extension can potentially reveal shared molecular signatures and cellular dynamics across various anti-aging interventions, providing valuable therapeutic targets for mitigating cell state and population changes in aging and aging-related neurodegenerative disorders.

## Methods

### Lead contact

Further information and requests for resources and reagents should be directed to and will be fulfilled by the lead contact, Junyue Cao.

### Materials availability

This study did not generate new unique reagents.

### Data and code availability

*EasySci* single-nucleus RNA seq data and *IRISeq* spatial transcriptomic data have been deposited at GEO and are publicly available as of the date of publication (accession number: GSE273038). Accession numbers are listed in the key resources table. All raw sequencing reads and processed files are accessible at BioProject PRJNA1138407.

The computational processing pipeline for processing *EasySci* data, including read alignment and gene count matrix generation is available on GitHub https://github.com/JunyueCaoLab/EasySci. The computational scripts for differential abundance and differential gene expression analysis are available on GitHub at this repository: https://github.com/zhangzehao626/PanSci. Any additional information required to reanalyze the data reported in this paper is available from the lead contact upon request.

### Experimental model and study participant details

#### Animals and sample collections

C57BL/6JN wild-type male mice brain samples were provided from the National Institute of Aging colony at Charles River. Caloric-restricted (CR) animals and *ad libitum* (AL) animals were housed separately and individually to monitor diet intake. Mice were weaned at 3-4 weeks of age and then caged in lots of four mice until 14 weeks of age. Caloric restriction begins at 14 weeks of age when animals are fed 90% of the predetermined amount of food consumed by AL age-matched controls. At 15 weeks of age, animals are fed 75% of the predetermined diet, and at 16 weeks are fed 60% of the predetermined diet. Metadata for each animal, including mouse individual ID, gender, age, accession dates, and body and brain weights, can be found in Table S1.

#### Whole Brain Nuclei Extraction for *EasySci*

The samples were processed by EasySci^10^ with minor modifications. In brief, frozen brain tissues were added to a 6 cm cell culture dish containing 3mL of lysis buffer solution (EZ Lysis Buffer containing 1% diethylpyrocarbonate). Each sample was homogenized with a razor blade until the solution could be aspirated with 1000uL tips to pass through a 40μM filter above a 50mL tube that contained 6 mL of lysis buffer solution. We then used the plunger from a 10mL syringe to further homogenize the tissue on the filter until the solution passed through the filter completely. The filters were washed with 1 mL of additional lysis solution. The nuclei were concentrated by centrifugation at 500g for 5 minutes at 4°C. The nuclei were washed 3 times in nuclei wash buffer (NWB), containing Nuclei buffer (NB, 10 mM Tris-HCl pH 7.5, 10 mM NaCl, 3 mM MgCl2 in RNase-free water) supplemented by 1% of 10% Tween-20 diluted in RNase-free water, 1% recombinant albumin (NEB, #B9200S), and 0.1% SUPERase•In™ RNase Inhibitor. After the washes, nuclei were aliquoted into 2 cryovials containing 500μL of nuclei resuspended in NWB containing 10% dimethyl sulfoxide and stored in slow freezers, cooling at 1°C per minute, at −80C overnight.

The following day, one aliquot of the nuclei from each sample was rapidly thawed in a 37 °C water bath and centrifuged at 500g for 5 minutes at 4°C. The supernatant was removed, and the nuclei were resuspended in freshly prepared NWB containing 0.005 mg/mL DAPI (Thermo Fisher, #D1306) for fluorescence-activated cell sorting (FACS). The nuclei were stained for 10 minutes before performing FACS on a SH800 Cell Sorter with a 100 μm sorting chip (Sony, #LE-C3210), to capture all DAPI-positive singlet nuclei and to exclude cellular debris and doublet cell populations. Nuclei were collected into a 1.5 mL DNA Lo-Bind tube containing 100 μL of NWB, vortexed to coat the tubes for optimal nuclei capture, and subsequently centrifuged at 500g for 5 minutes at 4 °C before proceeding directly to library construction.

#### *EasySci* library construction and sequencing

The subsequent steps for the generation of sequencing libraries followed the published *EasySciRNA* protocol (2). Initially, the sorted nuclei were distributed across 96-well plates (Geneseesci, #24-302) for reverse transcription (RT). Prior to RT, indexed oligo-dT and indexed random hexamer primers were utilized to introduce the first index. The nuclei were pooled, washed, and redistributed into new 96-well plates for the addition of the second index through ligation. After subsequent pooling and washing steps, the nuclei were diluted to 500 nuclei/uL and distributed into new plates for second-strand synthesis and 1x AMPure XP SPRI purification (Beckman Coulter, #A63882). After elution of the cDNA, tagmentation with Tn5 transposase was performed, followed by a 16-cycle PCR reaction. The resulting PCR products were then pooled and purified twice using 0.8X volume of AMPure XP SPRI. The library concentration and fragment size were measured using an Agilent TapeStation, and sequencing was carried out on an Illumina NovaSeq 6000 System with an S4 Flow Cell.

#### Sequencing data preprocessing

BCL files were demultiplexed using the barcode information from the last round of PCR indexing utilizing Illumina’s bclfastq2 program (version 2.20.0.422) to convert the file format to FASTQ. For single-nucleus RNA-seq, read alignment and gene/exon count matrix generation were conducted using our *EasySci* pipeline (https://github.com/JunyueCaoLab/EasySci). Cells from the gene count matrix were filtered according to the following parameters: unmatched rate less than 0.4, combined shortdT and randomN UMI (Unique Molecular Identifier) count less than 200, and gene counts less than 100. Scrublet (version 0.2.3) was utilized for the doublet removal set with the following parameters min_count = 3, min_cells = 3, vscore_percentile = 85, n_pc = 30, expected_doublet_rate = 0.08, sim_doublet_ratio = 2, n_neighbors = 30. Cells with a doublet score of more than 0.2 were discarded, which corresponded to 10% of the processed and filtered dataset.

#### Integration and label transfer

To increase the resolution of main cell population clustering, we integrated this dataset with the published EasySci brain atlas^10^. The EasySci brain atlas^10^ containing ∼1.5 million cells is firstly sampled to 126,285 cells by subsetting 5,000 cells from each main cell population (for cell populations that have less than 5,000 cells, we subset all cells). The subsampled EasySci brain atlas is then integrated with the full brain CR dataset using Seurat^44^ integration functions. Briefly, each dataset is normalized and the top 5,000 highly variable features are selected using “vst” method. The integration features are selected by SelectIntegrationFeatures() and anchors are determined using FindIntegrationAnchors(). The two datasets were then integrated with the IntegrateData() function. To visualize all the cells together, we co-embedded all the cells in the same low-dimensional space.

To transfer labels from the EasySci dataset to the brain CR dataset, we implemented a two-step label transfer process targeting both the main cell population and subcluster levels. First, we identified the common genes shared by both datasets and subsetted the datasets based on this shared gene set. The annotated EasySci dataset was then randomly subsampled to 1,000 cells per main cell population, retaining all cells if a cell population contained fewer than 1,000 cells. We employed a support vector machine (SVM) classifier, specifically the LinearSVC function from scikit-learn, to train the model using the log-transformed expression values of the subsampled cells, with the corresponding main cell populations serving as the target variable. The trained SVM model was evaluated by comparing it with a model trained on permutated data and was then applied to the brain CR dataset to predict the main cell population labels. For subcluster prediction, a similar approach was employed: for each EasySci main cell population, a maximum of 1,000 cells per subcluster was subsampled, and an SVM model was trained for each main cell population. Each classifier was then applied to the brain CR data, subsetted by the predicted main cell population labels, to predict the subcluster identities.

#### Differential gene analysis for single-nucleus data and spatial genomics

To identify differentially expressed genes for each main cell population between adult and aged groups and for each brain region between caloric restriction and *ad libitum* groups, we employed the likelihood ratio test to identify genes significantly associated with specific cell populations/subpopulations, using differentialGeneTest() function in Monocle2 (version 2.28.0). To ensure both statistical significance and biological relevance, we applied stringent selection criteria: 1) for identifying aging-associated genes (AAGs), we use the FDR threshold of 1e-5, the enrichment fold of a gene in the most highly expressed cell cluster versus the second highest is greater than 2, the maximum transcripts per millions (TPMs) over than 50 in specific conditions, and the cell number of a cell type is over 1,000 is *ad libitum* dataset (avoiding the bias introduced by low-aboundance cell type); 2) for identifying brain region-specific DE upon CR, we use the FDR threshold of 0.05, the enrichment fold of a gene in the most highly expressed cell cluster versus the second highest is greater than 1.5, and the maximum transcripts per millions (TPMs) over than 50 in specific conditions.

#### Cell state transition analysis by integrating datasets across studies

To identify detailed subpopulations of microglia, oligodendrocytes, and those involved in neurogenesis, we integrated corresponding main cell populations from this dataset, EasySci^10^, and TrackerSci^19^ (for identifying the potential progenitor states). Specific main cell populations were subset from each dataset and integrated using Seurat^44^ integration functions. Similar to the abovementioned full dataset integration, each dataset is normalized and the top 5,000 highly variable features are selected using “vst” method. The integration features are selected by SelectIntegrationFeatures() and anchors are determined using FindIntegrationAnchors(). The datasets were then integrated with the IntegrateData() function and co-embedded in the 2D space.

#### Differential abundance analysis for single-nucleus dataset

To assess cell population dynamics across different age groups and diet conditions, we conducted the differential abundance test as demonstrated in^42^. Briefly, cell count matrices across replicates (cell population X replicates) are constructed via counting cell numbers of each cell population and then normalized against the total cell number recovered from specific replicate. We then employed the likelihood-ratio test for identifying differentially abundant cell populations using the differentialGeneTest() function of Monocle2 (version 2.28.0). For fold change calculations, we first normalized the number of cells in each cell population relative to the total cell count in the respective condition. We then compared these normalized values between the case and control conditions, incorporating a small numerical value (10^−6^) to reduce the noise from very small clusters. To assess the significance of cell population dynamics of specific cell populations, we use an FDR threshold of 0.05 and a fold change higher than 1.5 between conditions.

#### Senescence score analysis

Gene expression counts in cells from each individual mouse and main cell type were pseudobulked by sum using the *decoupler-py* package. Groupings with fewer than. This data was normalized to 10,000 total counts and logarithmized with a pseudocount of 1; the Scanpy *score_genes* function was then used to calculate SenMayo score using the SenMayo mouse gene list downloaded from Saul *et al* (2022). Before plotting, pseudobulk groupings with fewer than 20 cells or 1000 counts were removed. Main cell types with fewer than 30 remaining mice were filtered out. For subcluster-specific score calculation, the same method was employed except that cells were pseudobulked by individual mouse and predicted subcluster; pseudobulk groupings with fewer than 10 cells or 1000 counts were removed.

To calculate SenMayo rescue significance, we used the edgeR package to fit pseudobulked count data (not normalized) to a generalized linear model, with age in months as a covariate and diet as a binary factor. For each main cell type, genes with fewer than 100 counts were removed; main cell types represented in fewer than 25 mice were also removed. The ROAST rotation gene set test^45^ was then performed using the SenMayo gene set for each cell type, with 1000 rotations. One-sided *p*-values were calculated for age and diet and multiple hypothesis correction was performed using the Benjamini-Hochberg method^45^.

#### Gene level analysis for neurogenic cells

Neurogenic cells (OB neurons 1_11 and OB neurons 1_17) were aggregated together and pseudobulked by sum for each individual 12-month-old mouse using the decoupler package. Mice with fewer than 10 cells or fewer than 100 expression counts were removed. Total counts per mouse were normalized and genes with fewer than 40 counts per million were removed. The differentialGeneTest() function from monocle2 was used to calculate significance of differential expression between 12-month-old AL mice and 12-month-old CR mice for each gene. Upregulated genes in CR were defined as having a *q*-value < 0.05 and a log-fold-change >= 1; downregulated genes in CR were defined as having a *q*-value <0.05 and a log-fold-change <= −1. Overrepresentation analysis was performed using Enrichr^46,47^ to identify biological processes from the GO Biological Process 2023 dataset that were significantly enriched in upregulated or downregulated genes.

#### *IRISeq* library construction and sequencing

To capture RNA from tissues, a beads array is generated by mixing ∼5 ul of receiver beads and sender beads in 6XSSC buffer in a 1:3 ratio, respectively. Then gel beads are monolayered into a square shape utilizing a gasket barrier of the desired shape and size, and semi-dried for a few minutes until their stabilization on the glass is observed, and no movement is observed. Then, 10-um thick tissue sections are placed on the beads array, and RNA hybridization is performed for ∼13 minutes without additional buffer addition to prevent array disturbance. Next, photocleavage of sender beads is performed for ∼2 minutes in a UV device (alpha thera AT8001). Library preparation follows protocols^48^, with slight modifications. Following second strand synthesis, cDNA was released from receiver beads utilizing 0.1 N NaOH and quenched with 1 M Tris-HCl pH 7.0, followed by 1X Ampure beads purification following manufacturer recommendations. cDNA amplification followed protocols^48^ utilizing SMRT primer and Truseq_Read1. Amplified cDNA tagmentation is adapted from (21). For receiver beads-sender beads DNA interactions library preparation, after collecting supernatant for cDNA library preparation, beads were washed with TE-TW buffer adapted from protocols^48^, and then resuspended in ∼25 ul of H2O. Then, DNA connections between sender and receiver beads were amplified by performing on-bead PCR and utilizing the NEBnext master mix. 0.25 uM of P5_Truseq_Read1 primers and Truseq_Read2 indexed sequencing primers were utilized for PCR. After PCR, perform gel extraction of the connections band which should be approximately ∼250 bp in size.

#### Spatial data processing and spatial map reconstruction

To reconstruct the spatial location of the bead array, interaction counts between sender and receiver beads were filtered to remove low-quality interactions, applying a cutoff of approximately 10 counts as the minimum threshold for considering a connection as real. From the filtered data, a matrix of sender-by-receiver bead counts was generated, then scaled and log1P transformed. This matrix was then subjected to PCA transformation, followed by UMAP analysis on a GPU using cuML, with parameters slightly adjusted from the *IRISeq* data^11^ to recreate the square-shaped array.

## Acknowledgments

We thank members of the Cao lab for helpful discussions and feedback. This research was made possible in part using biomaterials from the NIA Aged Rodent Tissue Bank (Aged Rodent Tissue Bank | National Institute on Aging (nih.gov)) at the University of Washington, Seattle under a contractual agreement with the National Institute on Aging (NIA). This work was funded by grants from the NIH (1DP2HG012522, 1R01AG076932, and RM1HG011014 to J.C). This work was supported by the Stavros Niarchos Foundation (SNF) and Kellen Women’s Entrepreneurship Fund at The Rockefeller University to W.Z., Diana Jacobs Kalman/AFAR Scholarships for Research in the Biology of Aging to Z.Z., and Medical Scientist Training Program grant from the National Institute of General Medical Sciences of the National Institutes of Health under award number: T32GM152349 to the Weill Cornell/Rockefeller/Sloan Kettering Tri-Institutional MD-PhD Program for A.A..

## Author contributions

W.Z. and J.C. conceptualized and supervised the project. C.S. and A.E. collected tissues and performed the *EasySci* experiment. Z.Z. and C.S. processed raw *EasySci* sequencing data. A.A. and W.J. performed the *IRISeq* experiment. A.A. processed raw *IRISeq* sequencing data. Z.Z. and A.E. performed the downstream analysis and generated all figures. W.Z., J.C, Z.Z. and A.E. wrote the manuscript with insights from all co-authors.

## Supplemental information

**Figure S1.**
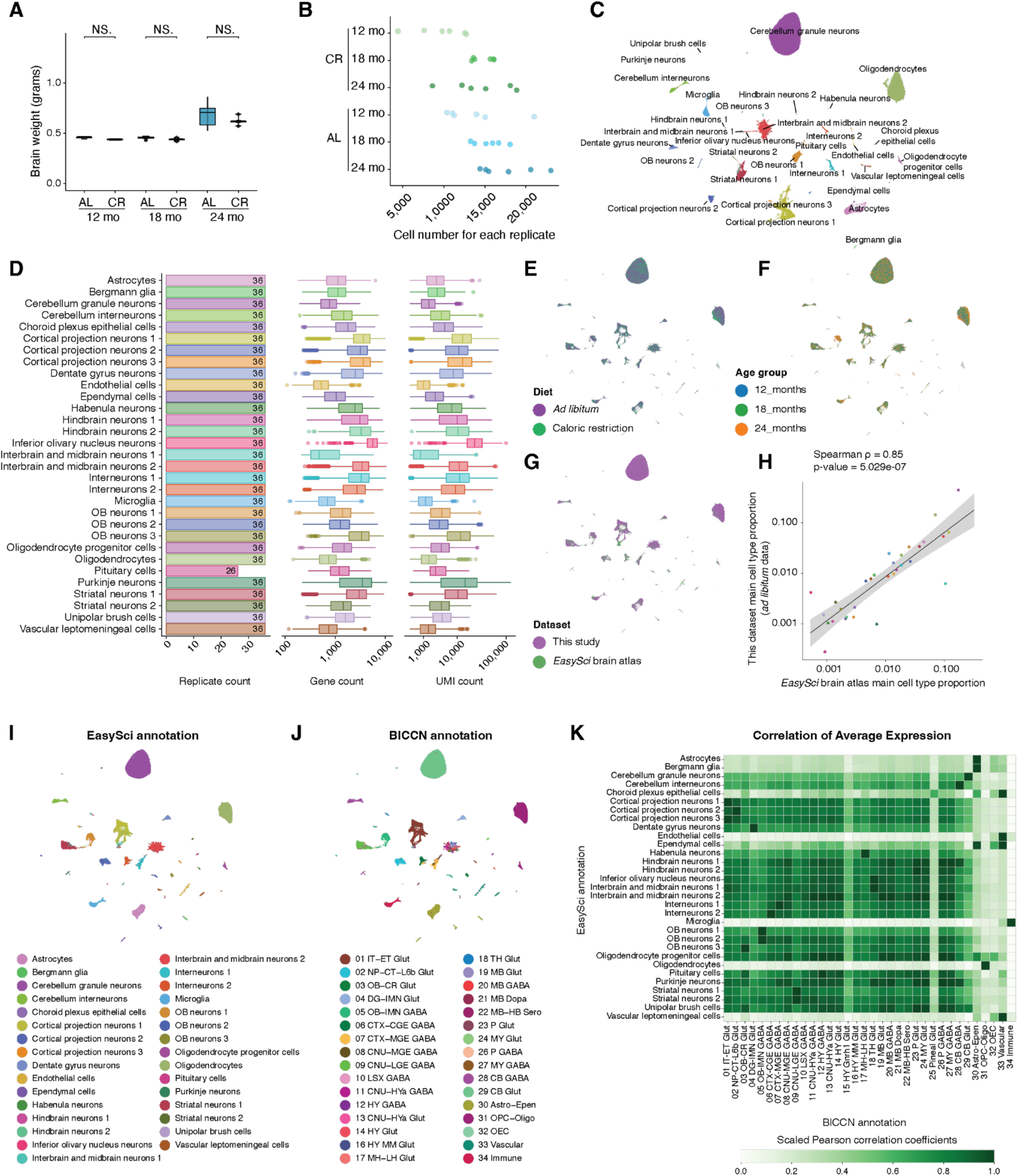
Performance analysis of the *EasySci* transcriptome profiling of mouse brains across various ages and in response to CR. **(A)** Brain weights across experimental groups. *Welch’s t-test,* ‘NS.’ represents not significant. **(B)** Scatter plots showing the number of cells recovered from each replicate in each experimental condition. **(C)** UMAP visualization of mouse brain cells from this CR brain atlas study, colored by main cell populations. **(D)** Bar plot (Left) and box plot (Right) showing the replicate number, gene number, and UMI number for cells of each main cell population. **(E)** UMAP visualization of mouse brain cells colored by diet conditions. UMAP coordinates were calculated using integrated data from both datasets (reference brain atlas^10^ and CR brain atlas). **(F)** UMAP visualization of mouse brain cells colored by age group. UMAP coordinates were calculated using integrated data from both datasets (reference brain atlas^10^ and CR brain atlas). **(G)** UMAP visualization of mouse brain cells integrating data from both datasets (reference brain atlas^10^ and CR brain atlas), colored by cells recovered from different studies. UMAP coordinates were calculated using integrated data consisting of the reference brain atlas^10^ and the CR brain atlas. Of note, the plot shows all cells from this study and sub-sampled data (Up to 1,000 cells per main cell population) from the published study^10^ used for integration analysis. **(H)** Scatter plot showing the fraction of all cells that each main cell type represents in the global brain population recovered by reference brain atlas^10^ (x-axis) or in this study (y-axis) **(I-J)** UMAP visualization of CR brain atlas, colored by main cell type annotation transferred from (I) EasySci or (J) Allen BICCN atlas. **(K)** Heatmap showing the min-max scaled Pearson correlation coefficients between the average gene expression profiles of EasySci-annotated cell types (rows) and BICCN-annotated cell types (columns). Higher values indicate stronger transcriptomic similarity between matched populations. Color bar denotes scaled correlation coefficients. Astro, astrocyte; CB, cerebellum; CGE, caudal ganglionic eminence; CNU, cerebral nuclei; CR, Cajal–Retzius; CT, corticothalamic; CTX, cerebral cortex; DG, dentate gyrus; Epen, ependymal; ET, extratelencephalic; HB, hindbrain; HY, hypothalamus; HYa, anterior hypothalamic; IMN, immature neurons; IT, intratelencephalic; L6b, layer 6b; LGE, lateral ganglionic eminence; LH, lateral habenula; LSX, lateral septal complex; MB, midbrain; MGE, medial ganglionic eminence; MH, medial habenula; MM, medial mammillary nucleus; MY, medulla; NP, near-projecting; OB, olfactory bulb; OEC, olfactory ensheathing cells; Oligo, oligodendrocytes; OPC, oligodendrocyte precursor cells; P, pons; TH, thalamus. Neurotransmitter types: Dopa, dopaminergic; GABA, GABAergic; Glut, glutamatergic; Sero, serotonergic.

**Figure S2.**
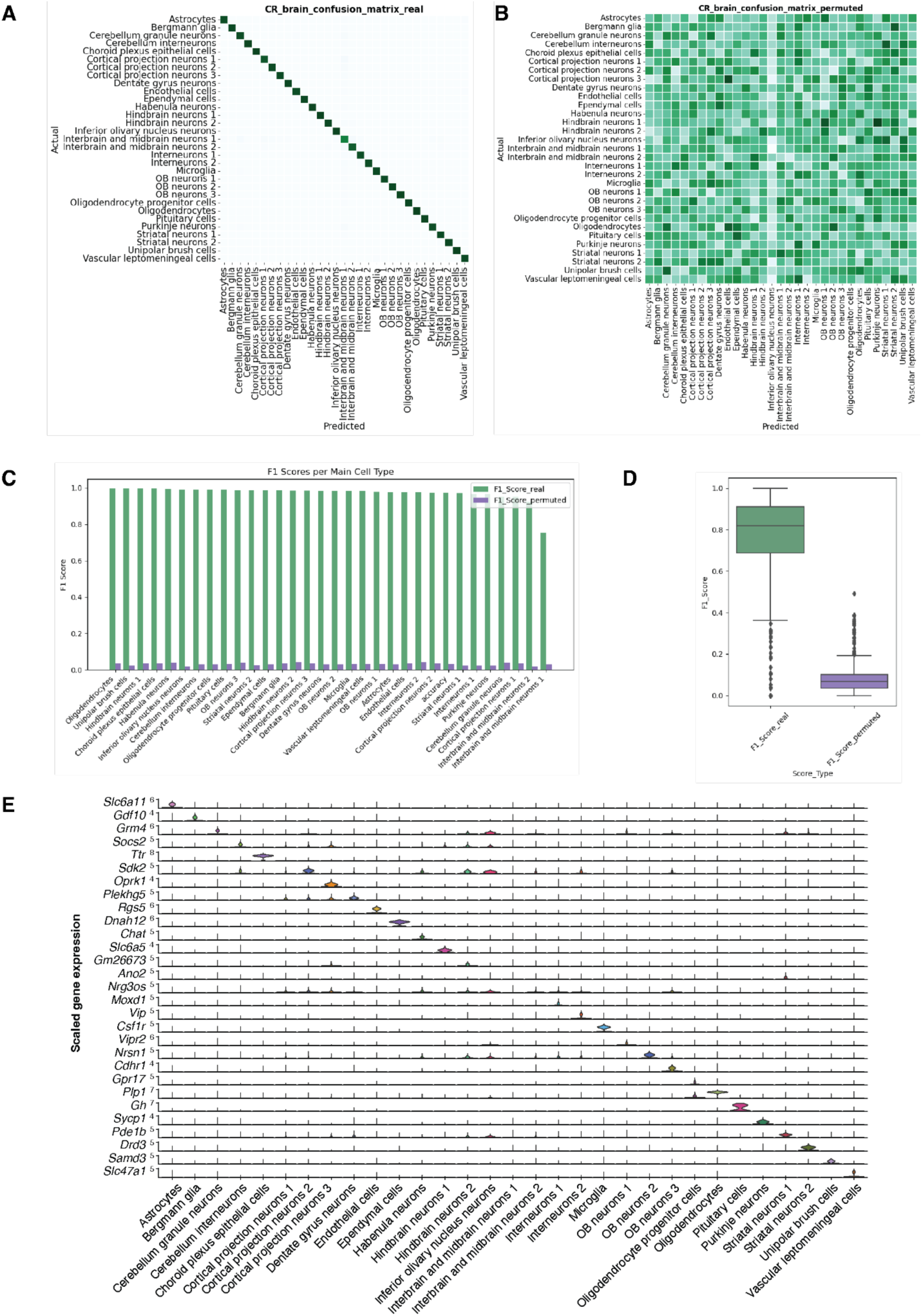
Support vector machine cross-validation performance for main cell type and subtype-level cell type annotation. (A) Confusion matrix showing predicted versus actual labels for main cell types using 5-fold cross-validation on the reference dataset. (B) Confusion matrix for permuted labels, serving as a baseline control. (C) Barplot comparing the F1 scores comparing real (green) versus permuted (purple) labels across all main cell types. (D) Boxplot showing F1 score across all subclusters, comparing real (green) versus permutated (purple) labels. **(E)** Violin plot showing the gene expression specificity across main cell populations, with expression data aggregated, normalized, and scaled for each main cell population.

**Figure S3.**
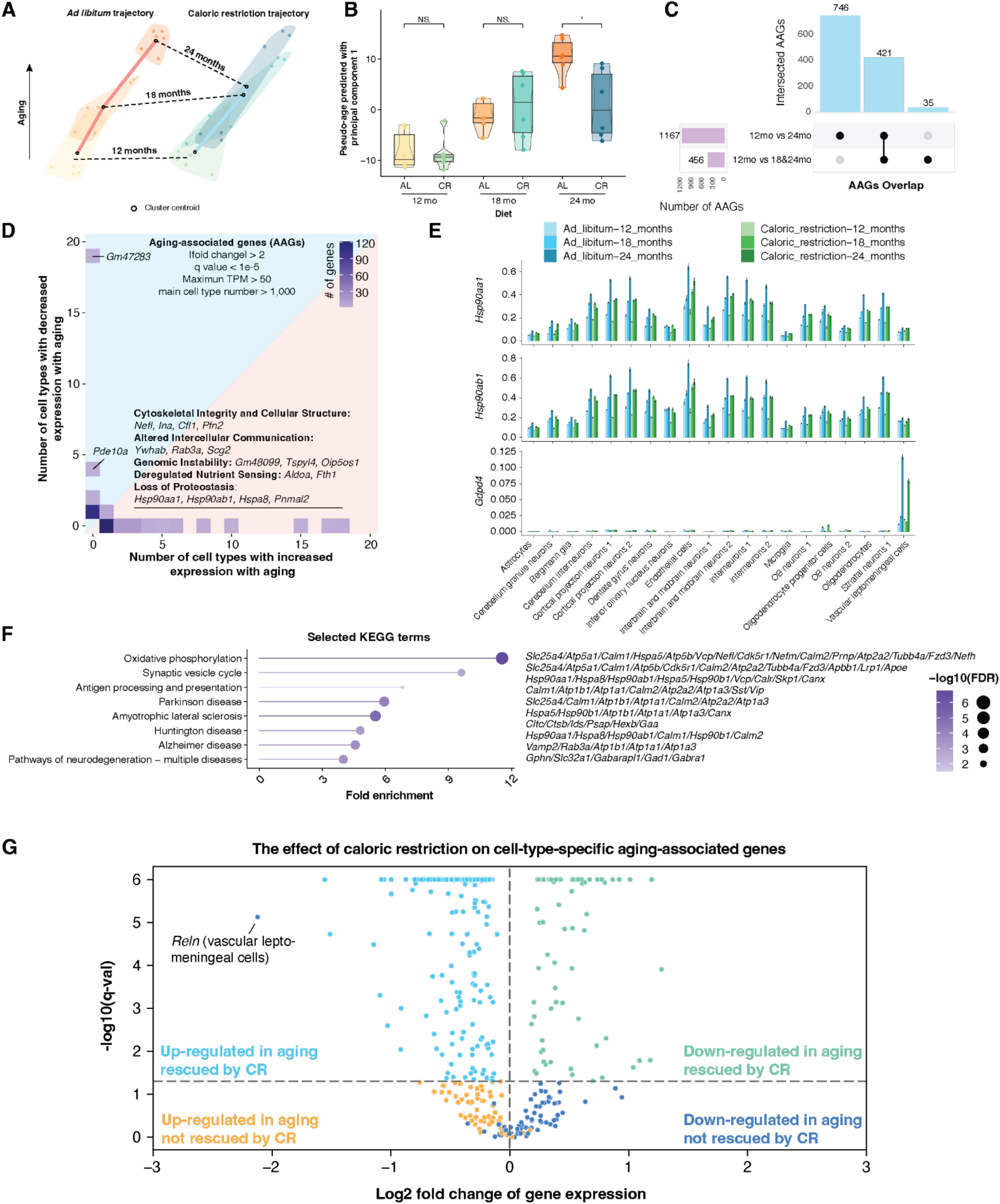
Identification of cell-type-specific gene expression changes in aging and upon CR. (A) Dimension reduction plot showing the pseudobulked transcriptomes aggregated from aging-associated genes for each replicate. The color indicates replicates from each age group, and the centroid of each age group is labeled. (B) Box plot comparing the principal component 1 score, reflecting the aging trajectory, for each condition. Welch’s *t*-test,* represents p-value < 0.05, and ‘NS.’ represents not significant. (C) Upset plot depicting the intersection of aging-associated genes identified in comparisons between 12-month and 24-month groups, and between adult (12-month) and aged (combining 18-month & 24-month groups). AAGs were identified based on false discovery rate (FDR) < 1e-5, expression in the most highly expressed cell type at least twice that in the second highest, maximum transcripts per million (TPM) > 50, and originating from cell types with over 1,000 cells in the *ad libitum* dataset. **(D)** Heatmap displaying the number of cell populations in which each aging-associated gene either increases (x-axis) or decreases (y-axis) in expression during aging. The color gradient from light to dark blue indicates the number of genes, with darker shades representing higher numbers. Broadly altered aging-associated genes (AAGs) are highlighted in the text. **(E)** Barplots showing the scaled expression (with error bars representing the standard error) of genes upregulated with aging but rescued by caloric restriction, across main brain cell populations under *ad libitum* and caloric restriction conditions. **(F)** Similar to Figure 2G, a lollipop plot showing the top enriched KEGG terms with a list of increased aging-associated genes that are rescued by CR. Detailed genes are listed near the KEGG terms. **(G)** Volcano plot showing effect of caloric restriction on cell-type-specific aging-associated genes. All genes shown are significantly upregulated (cyan/orange) or downregulated (green/purple) in aging. X-axis represents the log2-fold-change in expression between 24mo AL mice and 24mo CR mice; Y-axis represents the significance of the expression change between 24mo AL mice and 24mo CR mice. The black dotted line represents the significance threshold for expression change between AL and CR. *Reln* (in vascular leptomeningeal cells) is highlighted as it is significantly downregulated in both aging and CR.

**Figure S4.**
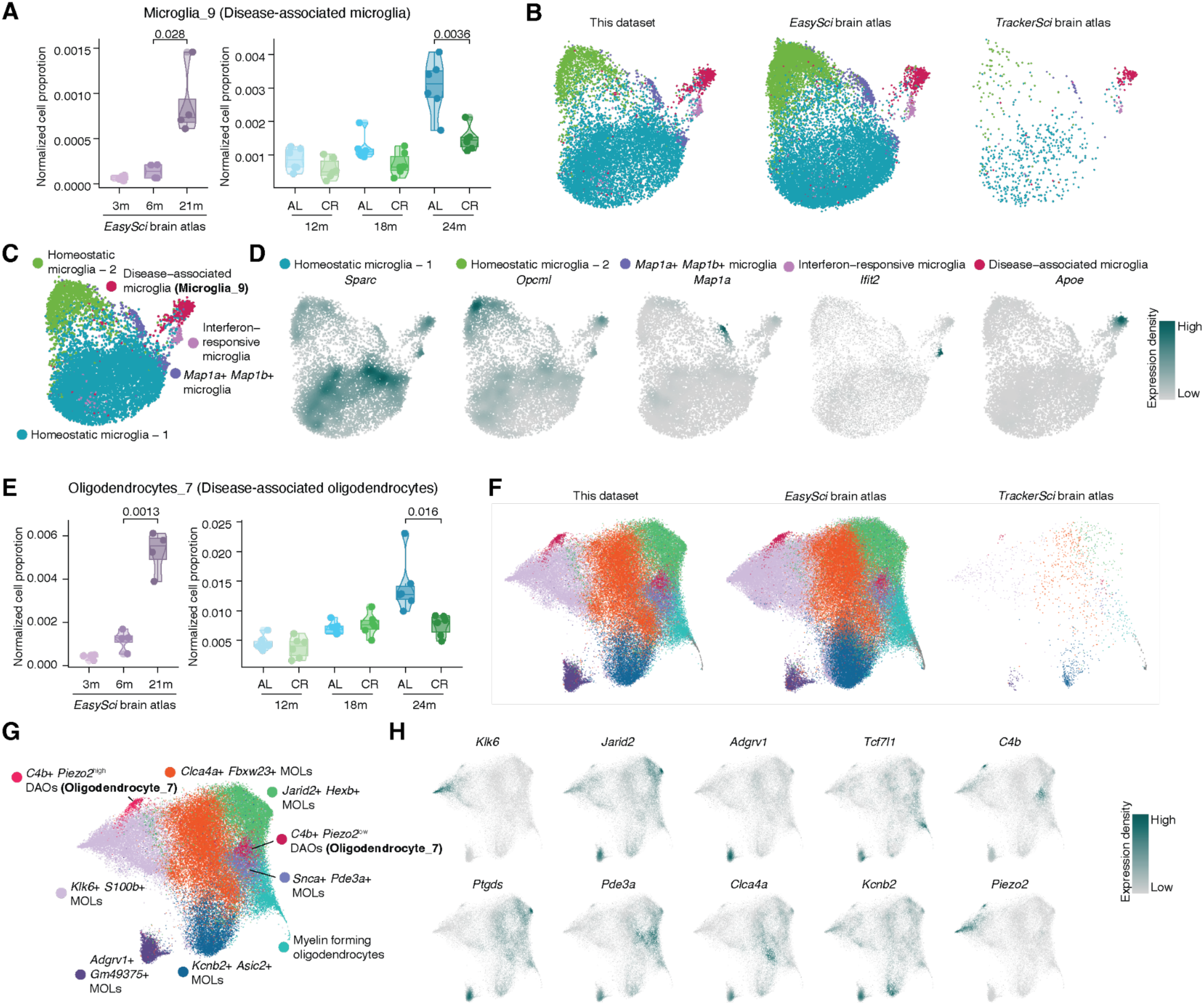
Identification of aging-associated subpopulations of microglia, oligodendrocytes, and neurogenic cells. **(A)** Box plot showing cell proportion changes in disease-associated microglia (DAM) across different ages and diet conditions in the brain aging atlas^10^ (left) and this dataset (right). **(B)** UMAP visualization of microglia by integrating cells from this study and two external datasets^10,19^, split by the data source and colored by subpopulation annotation. **(C-D)** UMAP visualization of microglia subpopulations, colored by subpopulation annotation (C) or the density of subpopulation-specific marker expression (D). **(E)** Box plot showing the cell proportion changes in disease-associated oligodendrocytes across different ages and diet conditions in the brain aging atlas^10^ (left) and this dataset (right). **(F)** UMAP visualization of oligodendrocytes by integrating cells from this study and two external datasets^10,19^, split by the dataset and colored by subpopulation annotation. **(G-H)** UMAP visualization of oligodendrocyte subpopulations, colored by subpopulation annotation (G) or the density of subpopulation-specific markers expression (H).

**Figure S5:**
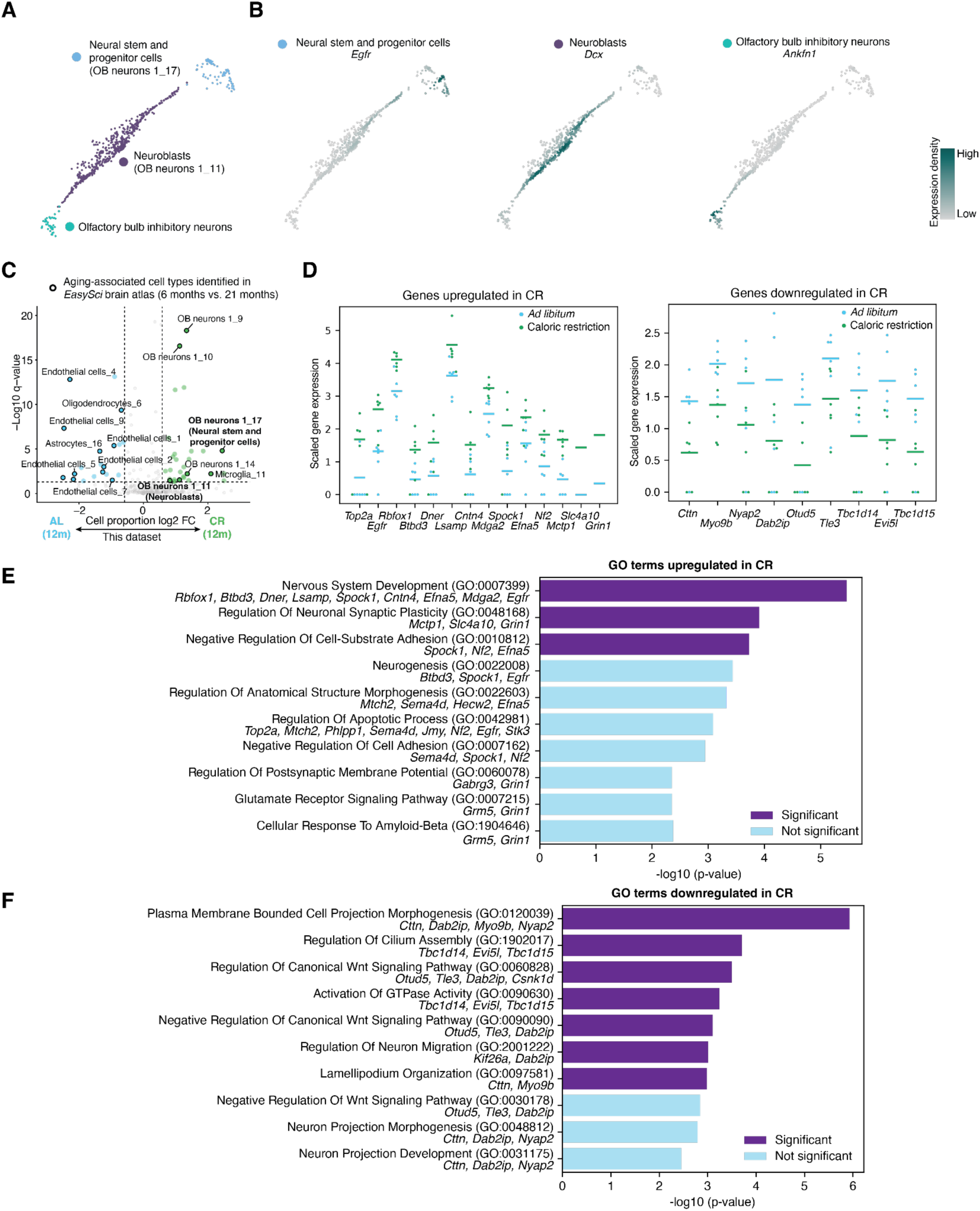
Effect of caloric restriction on olfactory bulb neurogenic cells in 12-month-old mice. **(A-B)** UMAP visualization of cell populations involved in neurogenesis, colored by subpopulation annotation (A) or the density of subpopulation-specific markers expression (B). **(C)** Volcano plot showing the cell subpopulation populations significantly altered between caloric restriction and *ad libitum* animals at 12 months. Aging-associated cell populations identified in the reference EasySci brain aging atlas^10^ were highlighted in a black circle. **(D)** Pseudobulk expression in 12-month mice of selected genes from these processes, which are upregulated or downregulated in CR. Each point represents the log-normalized expression in one mouse; horizontal lines represent the mean log-normalized expression across mice. *Top2a* is a common marker of cell proliferation. **(E-F)** GO Biological Processes (E) upregulated and (F) downregulated in CR relative to AL in 12mo mice.

**Figure S6.**
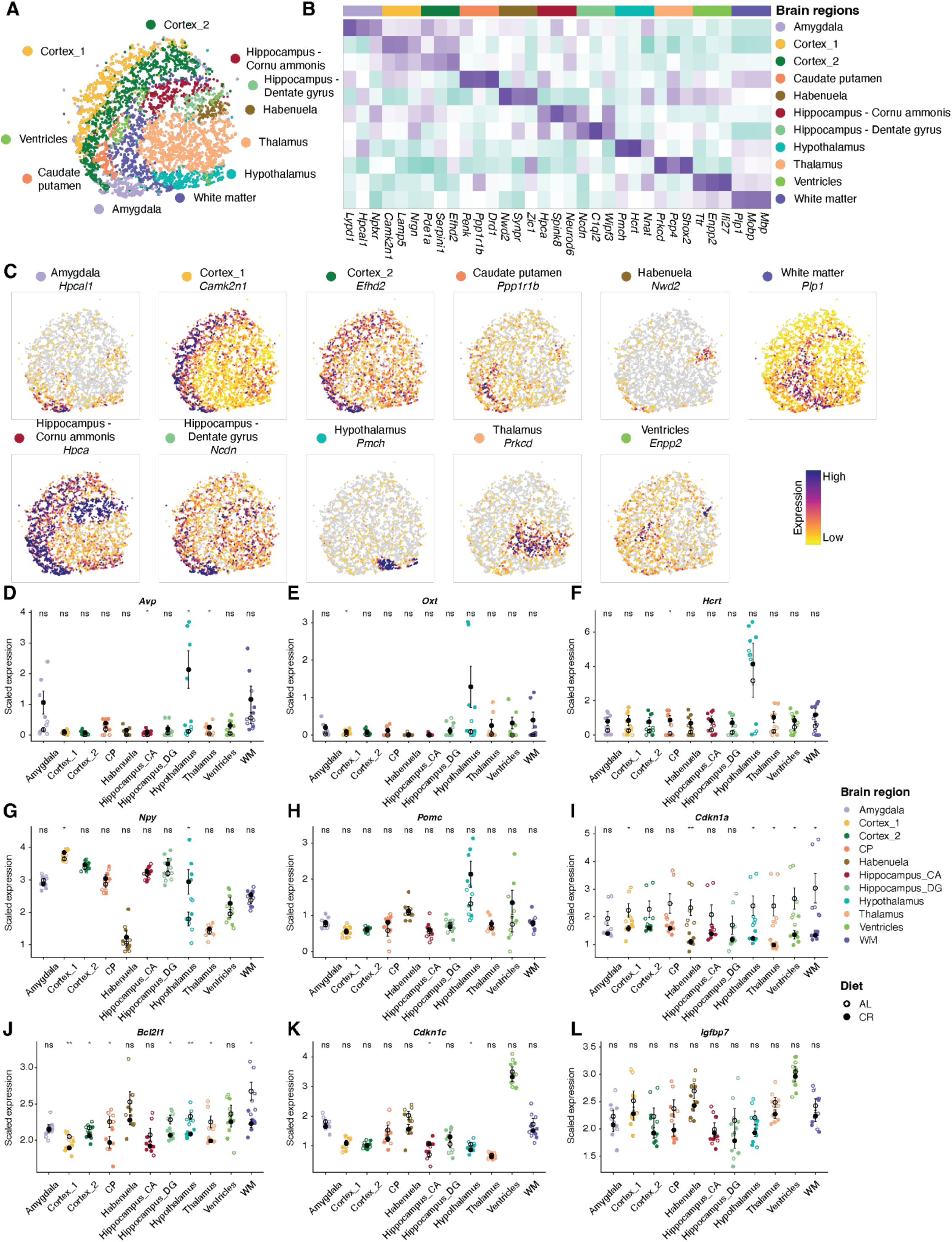
Identification of region-specific gene expression in the aged mouse brain sections. **(A)** Image reconstruction of all receiver beads derived from bead-bead connections, with each bead colored by annotated brain regions. **(B)** Heatmap showing the gene expression specificity across regions, with expression data aggregated, normalized, and scaled for each region. **(C)** Image reconstruction of all receiver beads derived from bead-bead connections, with coloring corresponding to the expression of region-specific gene markers. The raw gene expression count per spot is normalized by the total count, multiplied by 10,000, and log-transformed. (**D-L**) Dot plot showing aggregated expression for each replicate between 24-month AL and 24-month CR across brain regions for (D-H) hypothalamus neuropeptide *Avp*, *Oxt*, *Hcrt*, *Npy*, *Pomc,* and (I-L) senescence-associated genes *Cdkn1a, Bcl2l1, Cdkn1c, Igfbp7.* Gene expression of each replicate is compared between diet groups within each brain region using Welch’s *t*-test. ** represents p-value < 0.01, * represents p-value < 0.05, and ‘ns’ represents not significant.

**Figure S7.**
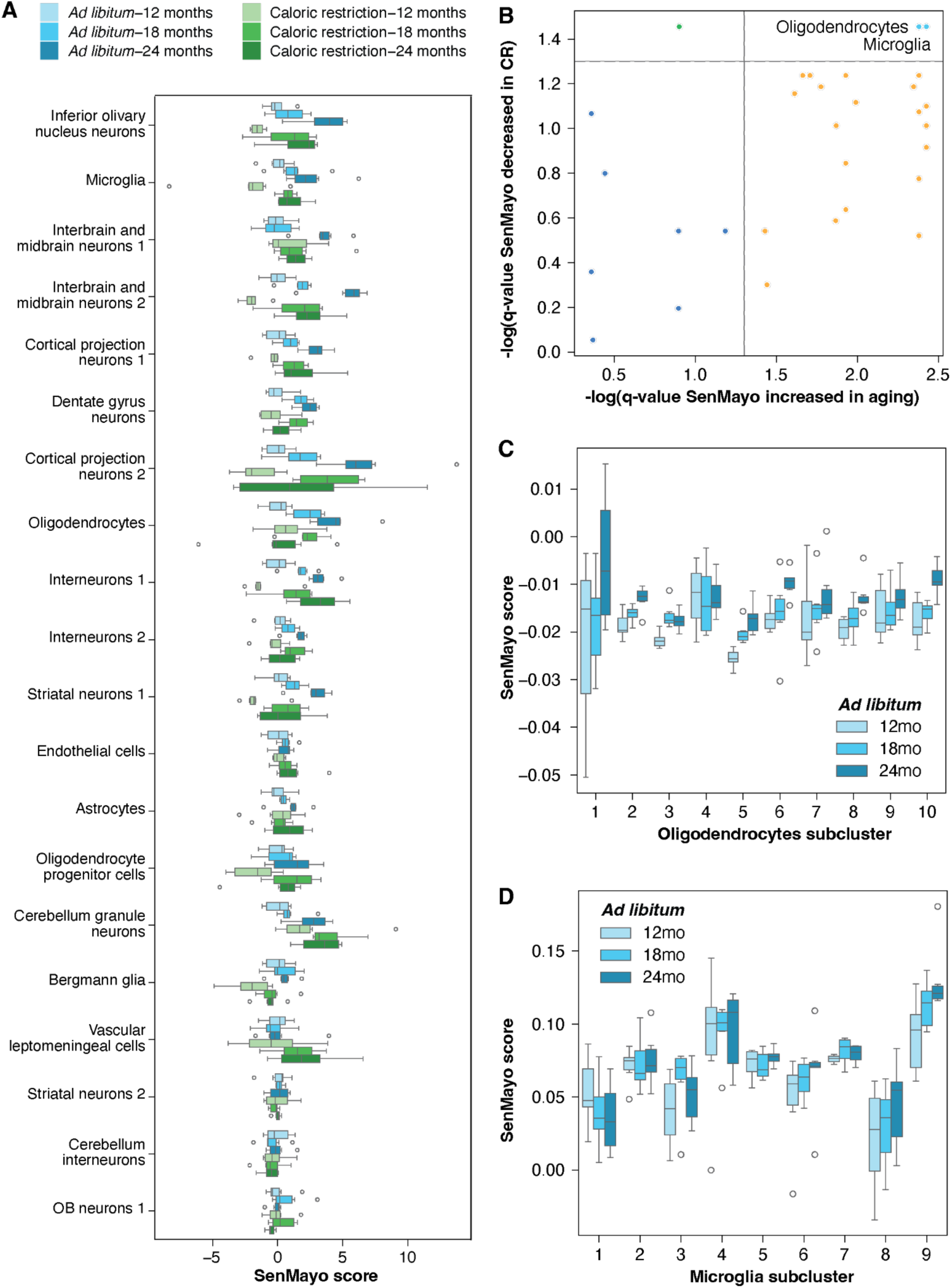
Caloric restriction rescues the aging-associated increase of senescence in glia cells. **(A)** Boxplot showing the effect of age and diet on SenMayo score in each main cell type (pseudobulked by individual mouse). For each cell type, the mean score in 12-month-old AL mice is subtracted from each score, and the result is divided by the standard deviation. Cell types are ordered by the magnitude of SenMayo change in aging AL mice. The center line represents the median; box spans 1st to 3rd quartiles. **(B)** Both oligodendrocytes and microglia exhibit an aging-associated increase of SenMayo score, which is rescued in CR. Each point represents one cell type; horizontal and vertical dashed lines represent significance thresholds. (**C-D**) Boxplots showing SenMayo score as a function of age in ad libitum oligodendrocyte (C) and microglial (D) cell subtypes. Center line represents median; box spans 1st to 3rd quartiles.

**Tables S1 to S3**

**Table S1. Metadata for mouse replicates included in the study.** For each animal and tissue section, this table describes the mouse replicate ID, sex, strain, age, diet group, body weight, brain weight, dissection date, and time and method used to profile this sample.

**Table S2. Aging-associated differentially expressed genes across main cell populations.** The “cluster_id” indicates which main cell populations this differentially expressed gene is proposed from. The “qval_aging”, “fold.change_aging”, and “class_aging” denote the FDR, fold change of the average expression between the adult and aged group in *ad libitum* condition, and the age group enriched for this gene. The “qval_CR”, “fold.change_CR”, and “class_CR” denote the FDR, fold change of the average expression between caloric restriction and *ad libitum* conditions in the aged group, and the dietary condition enriched for this gene. The “rescued_cellcount_type” shows if an AAG is shared by multiple cell populations or enriched in the specific cell populations. The “rescued_type” shows if an AAG is rescued by CR and the rescuing types.

**Table S3. Differentially expressed genes across brain regions upon caloric restriction.** For each gene, the “max.condition” is the condition with the highest expression (“max.expr”). The “second.condition” is the condition with the second highest expression (“second.expr”). The “fold.change” is the fold change between the max expression and second max expression. The “qval” is the false detection rate (one-sided likelihood ratio test with adjustment for multiple comparisons) for the gene differential expression test across different main cell populations. The “brain_region” denotes which brain region the DE gene is coming from.

## Notes

### Competing Interest Statement

The authors have declared no competing interest.

